# Sex specificity of root exudates and their effects on rhizosphere microorganisms in a rare and endangered plants, *Taxus cuspidata*

**DOI:** 10.64898/2026.10.14.759288

**Authors:** Wu Guoan, Zhang Yanwen

## Abstract

Dioecious plants exhibit sexual dimorphism not only in morphological structure, physiological metabolism, and resource acquisition but also in belowground microecosystems. In this study, high-performance liquid chromatography (HPLC) and high-resolution mass spectrometry (HRMS) were employed to detect sex-specific differences in root exudates of the typical dioecious plant *Taxus cuspidata* between adult and juvenile stages. High-throughput sequencing was used to analyze sex- and stage-specific differences in rhizosphere microbial communities. The results showed that in the adult stage, female plants enriched 2-mercaptoethanesulfonic acid and α-keto-γ-methylthiobutyric acid, while male plants secreted more terpenoids (e.g., α-humulene) and purine metabolic intermediates (xanthine). In the juvenile stage, female plants exhibited higher expression of phosphoenolpyruvate and L-valine, whereas male plants enriched 5-methylcytosine and tricyclic sesquiterpenoid antibiotics. The diversity of rhizosphere microorganisms showed a sex-reversal pattern across different developmental stages. Correlation analysis revealed that in the adult stage, the abundance and diversity of bacteria (Proteobacteria) and fungi (Mortierellomycota) were higher in female than in male plants, which were enriched with Actinobacteria. In the juvenile stage, male plants exhibited greater diversity of both rhizosphere bacteria and fungi than female plants, with enrichment of stress-tolerant and decomposing microorganisms (e.g., *Humicola*). The findings reveal different resource allocation and environmental adaptation strategies in the belowground parts of dioecious plants, which are of great value for the conservation of rare and endangered plants.

**Author summary:** Funding Source: This work was supported by National Natural Science Foundation of China (Grant No. 32272757)

The raw sequence data reported in this paper have been deposited in the Genome Sequence Archive (Genomics, Proteomics & Bioinformatics 2025) in National Genomics Data Center (Nucleic Acids Res 2025), China National Center for Bioinformation / Beijing Institute of Genomics, Chinese Academy of Sciences (GSA: CRA045178) that are publicly accessible at https://ngdc.cncb.ac.cn/gsa.

data link: https://ngdc.cncb.ac.cn/gsa/browse/CRA045178

## Introduction

The plant rhizosphere is a critical micro-zone where material exchange, energy flow, and information transfer occur between roots and the soil environment. Root exudates, collectively referring to various organic compounds released by plant roots into the soil, serve as key mediators of plant-soil-microbe interactions[1–3]. Root exudates mainly include primary and secondary metabolites as well as specific chemical signaling compounds[4–6]. They not only selectively recruit specific rhizosphere microorganisms[7–8], but also provide carbon sources, nitrogen sources, and other nutrients to rhizosphere microbes, and act as signaling molecules regulating microbial colonization, proliferation, and community structure. In turn, rhizosphere microorganisms influence plant growth, development, and stress tolerance by mobilizing soil nutrients, producing growth-promoting substances, and degrading toxic compounds, thereby forming a tightly interacting symbiotic system that jointly maintains the microecological balance of the plant rhizosphere [9,10].

Dioecy represents an important sexual differentiation type in plants. Due to differences in reproductive strategies, resource allocation, and metabolic pathways, male and female plants exhibit significant sexual dimorphism in morphological traits, physiological and biochemical characteristics, and stress tolerance[11–13]. In addition to aboveground parts, physiological activities in the belowground parts of dioecious plants also display sex-related differences. The composition, concentration, and secretion rate of root exudates show pronounced sex specificity[14–16]. This specificity not only directly alters the physicochemical properties and nutrient availability of the rhizosphere soil but also differentially regulates the community composition, diversity, and function of rhizosphere microorganisms [17,18].

Plant sex is an important factor shaping the structure of rhizosphere microbial communities. For example, root exudates of male and female poplar, papaya, and mulberry exhibit significant sex specificity[14,19–21], leading to the enrichment of different dominant microbial groups in the rhizosphere of each sex[22–25]. Male plants generally possess more complex bacterial co-occurrence networks[26], whereas female plants exhibit higher fungal abundance under specific environmental conditions[27]. Furthermore, sex specificity in root exudate composition is also associated with developmental stage. As plants grow and develop, root exudates often shift from being dominated by primary metabolites in juvenile stages to secondary metabolites in adult stages[28–30], thereby driving dynamic succession of rhizosphere microbial communities.

*Taxus cuspidata*, a species of the genus Taxus in the family Taxaceae, is a relict plant of the Tertiary glacial period and is primarily distributed in Northeast China and adjacent regions [31]. Due to its high ecological, medicinal, and scientific value, it has been listed as a National First-Class Key Protected Wild Plant in China. Although extensive research has been conducted on the genus Taxus, little is known about the characteristics of its belowground rhizosphere microecosystem. Specifically, it remains unclear whether sexual dimorphism also exists in the underground parts of this species, and whether such sex-specific differences undergo transitions at different developmental stages. Addressing these questions will help elucidate the effects of root exudates of dioecious plants on the structure, diversity, and function of rhizosphere microbial communities, thereby clarifying the interaction mechanisms among female/male plants, root exudates, and rhizosphere microorganisms.

## Materials and methods

### Study Site

The sampling site of this study was located on a hillside in Heigou Village, Jinshan Town, Dandong City, Liaoning Province (E123°62′, N40°08′). The site is a natural mountainous environment with a slope of approximately 20°, a soil layer thickness of 40 -50 cm, The soil at the sampling site is classified as brown soil, primarily consisting of sandy loam and light loam. Its chemical profile indicates a slightly acidic reaction, and is inhabited by *Taxus cuspidata* trees of various ages.

### Collection of Experimental Samples

Sampling was conducted in early May 2025 (when male plants had flowered and shed pollen, and female plants had visible female cones). Three male and three female plants of *Taxus cuspidata* growing in the same habitat were selected for each age group: 30-year-old adult plants and 6-year-old saplings, with individuals spaced approximately 5 m apart. A carbon-free mixed nutrient solution (main components: NH₄NO₃, KH₂ PO₄, K₂SO₄, MgSO₄, and CaCl₂) was pre-prepared as the root culture medium. For plants of different ages, the surface soil was first carefully removed to expose intact, undamaged living roots (diameter≤2 mm, 10-15 cm long). The roots were gently rinsed with purified water to remove adhering soil, then slowly transferred into a glass syringe with cotton placed at the bottom. Sterile glass sand was subsequently added to the syringe to completely cover the roots. Twenty milliliters of nutrient solution were injected into the device to rinse the roots and then withdrawn. Finally, another 20 mL of nutrient solution were added, and the roots were cultured for 24 hours. Root exudates were continuously collected for three days. The obtained samples were filtered thro ugh a 0.22 μm filter membrane and stored at -4°C for subsequent analysis.

Simultaneously with root exudate collection, rhizosphere soil tightly adhering to the roots of the sampled plants was collected at a depth of 10-15 cm. All samples were stored at -80°C for subsequent analyses.

### Analysis of Root Exudates

Root exudate samples were mixed with 0.4 mL ultrapure water, dissolved by ultrasonication, filtered through a microporous filter membrane, and transferred into autosampler vials. Data acquisition was performed using a Thermo Ultimate 3000 ultra-high-performance liquid chromatography system coupled with an Orbitrap Elite high-resolution mass spectrometer.

Chromatographic conditions: An Agilent SB-C18 column (1.8 μm, 2.1 mm×100 mm) was used at a flow rate of 0.35 mL/min, column temperature of 40°C, and injection volume of 4 μL. The mobile phase consisted of A: ultrapure water (with 0.1% formic acid) and B: acetonitrile (with 0.1% formic acid).

Gradient elution: 0-9 min, linear gradient from 5% to 95% B; 9-10 min, hold at 95% B; 10-14 min, maintain 5% B for equilibration.

Mass spectrometry conditions: Ion source temperature 300°C; ion source voltage: positive mode 3.5 kV / negative mode 2.5 kV; transfer tube temperature 275°C; sheath gas and auxiliary gas: 35 and 10, respectively; scan range 100-1500 m/z. Data-dependent acquisition (DDA) mode was used with MS² collision energy set to CID 35 eV.

Data analysis was performed using Compound Discoverer software for peak extraction, and compound identification was carried out by matching retention time (Rt), MS1, a nd MS2 information against local and public databases. Microbial Genomic DNA Extraction, PCR Amplification, and Sequencing

A 0.5-g aliquot of rhizosphere soil was mixed with extraction lysis buffer in a centrifuge tube and ground using a Tissuelyser-48 multi-tissue grinder at 60 Hz.

Primer selection: The V3-V4 region of the bacterial 16S rRNA gene was amplified using primers 338F/806R; fungi were amplified using primers targeting the ITS5 and IT S2 regions.

PCR amplification: Initial denaturation at 98°C for 3 min; 27 cycles (98°C for 30s, 5 5°C for 30s, 72°C for 45s); final extension at 72°C for 10 min; hold at 12°C.

Each sample was run in triplicate. PCR products were quantified using the Quant-iT PicoGreen dsDNA Assay Kit, pooled in equimolar ratios, and library construction was performed with the Illumina TruSeq Nano DNA LT Library Prep Kit. After quality control on an Agilent Bioanalyzer, sequencing was carried out on an Illumina NovaSeq 6000 platform using the NovaSeq 6000 SP Reagent Kit with 2×250 bp paired-end reads.

### Bioinformatics and Statistical Analysis

Data preprocessing: Raw sequences were processed using the DADA2 pipeline to rem ove primers, perform quality filtering, denoise, and merge reads, generating an amplicon sequence variant (ASV) table followed by chimera removal.

Taxonomic annotation: Bacterial 16S ASVs and fungal ITS ASVs were aligned against the SILVA (or Greengenes) and UNITE databases, respectively, to obtain taxonomic information.

Diversity analysis: After data normalization, alpha diversity indices (Chao1, Shannon, Simpson) were calculated. Beta diversity was analyzed using UniFrac distances, and differences in community structure among samples were visualized via principal coordinate analysis (PCoA) and non-metric multidimensional scaling (NMDS).

Differential significance testing: ANOVA with post-hoc tests (Dunn’s test, Steel-Dwass, etc.) was used to visualize differences in alpha diversity among sample groups. Permutational multivariate analysis of variance (PERMANOVA; Adonis) and analysis of similarities (ANOSIM) were employed to test for significant differences in community structure between groups. Linear discriminant analysis effect size (LEfSe) was used to identify significantly differentiated biomarkers.

To predict the correlations between root exudates and microorganisms, 20 substances with significant differences in root exudates were selected for Spearman correlation analysis with rhizosphere microbial communities at the genus level. The results are presented in Table 4 and Table 5.

## Results

### Sex Specificity of Root Exudates in Male and Female Plants

A total of 110 compounds were identified in the root exudates of *Taxus cuspidata*. The major categories of compounds and the proportions of compounds contained therein are shown in Table 1.

**Table 1.** Classification of root exudates.

| Compound Category | Number of Compounds | Proportion (%) |
| --- | --- | --- |
| Amino acids and their metabolic derivatives | 18 | 16.3% |
| Nucleosides, nucleotides and their bases | 8 | 7.2% |
| Vitamins and coenzymes | 7 | 6.3% |
| Lipids, fatty acids and their derivatives | 13 | 11.8% |
| Carbohydrates and their derivatives | 4 | 3.6% |
| Phenols, aromatics and organic acids | 19 | 17.2% |
| Terpenoids and steroids | 12 | 10.9% |
| Nitrogen-containing heterocycles, alkaloids and synthetic intermediates | 26 | 23.6% |

### Sex Specificity of Root Exudates in Adult Plants

Comparison of the top 10 root exudate compounds with the greatest differences between female and male plants showed that 9-Methyl-3-tropinone and 3-amino-2,3-dihydroxybenzoic acid were unique to female plants. α-Keto-γ-methylthiobutyric acid (female/male ratio 10.4) and 2-mercaptoethanesulfonic acid (female/male ratio 5.1) showed significant differences between sexes, while cinnamic acid (female/male ratio 9.8) is a major intermediate in the phenylpropanoid metabolic pathway (Fig 1a and 2a).

**Fig 1.**
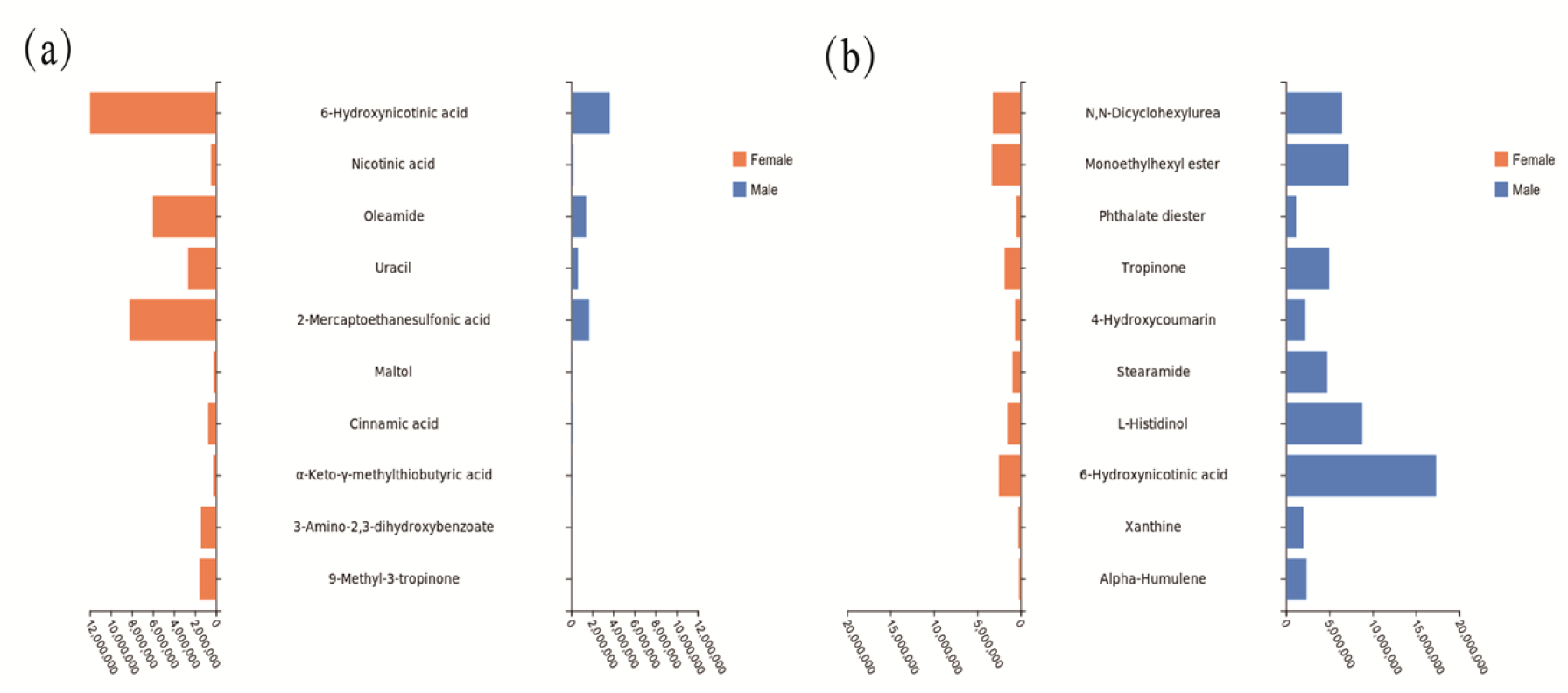
Pyramid plot of differential peak areas of root exudates in Taxus cuspidata. (a) represents the peak areas of the top 10 substances with higher peak areas in female plants than in male plants. (b) represents the peak areas of the top 10 substances with higher peak areas in male plants than in female plants.

**Fig 2.**
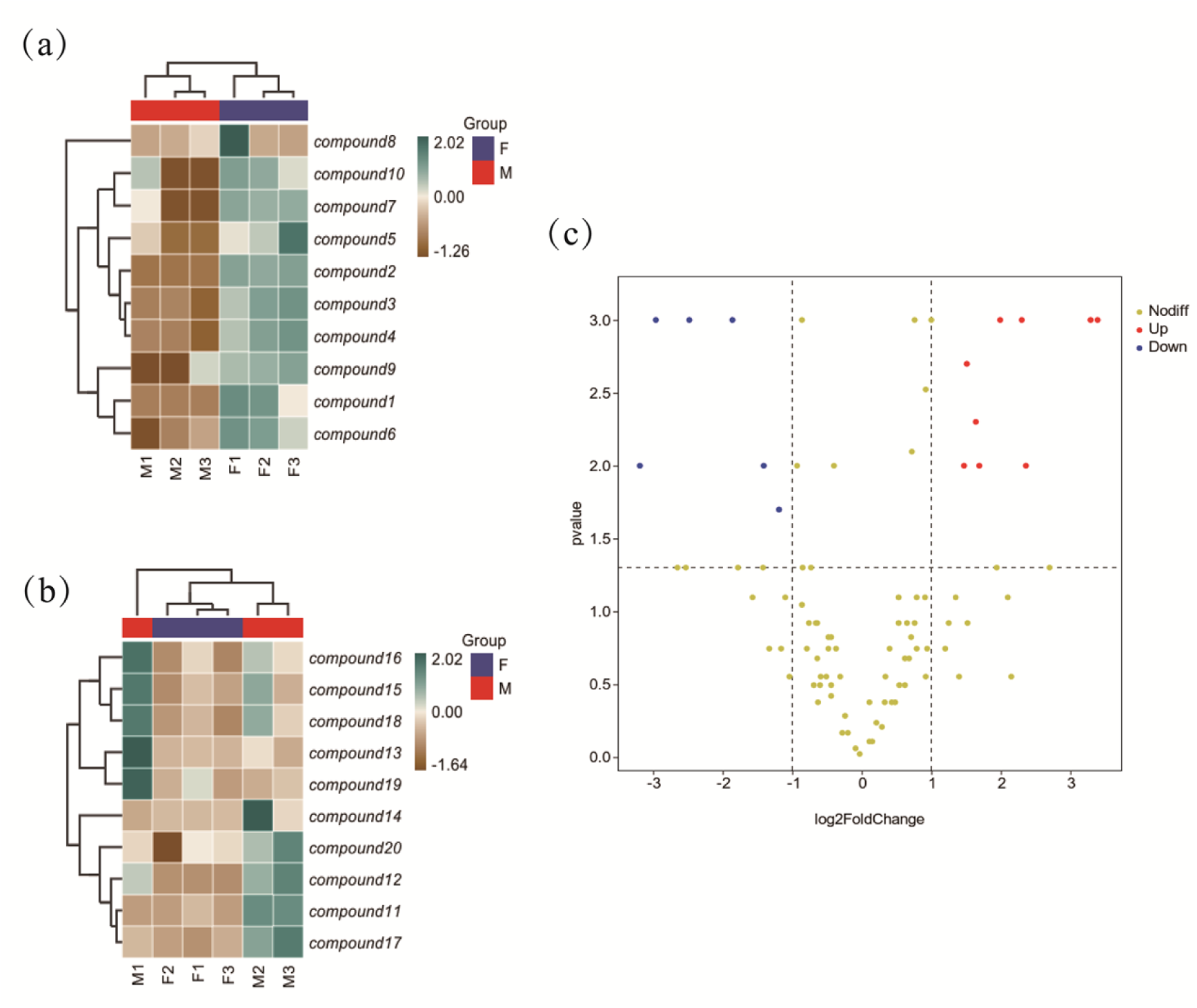
Heatmap and volcano plot of differential root exudates in Taxus cuspidata. (a) represents the heatmap of the top 10 substances with higher abundance in female plants than in male plants. (b) represents the heatmap of the top 10 substances with higher abundance in male plants than in female plants. (c) represents the volcano plot of significant differences in root exudates between male and female plants.

Among the compounds with higher abundance in male plants than in female plants, A LPHA-humulene showed the most pronounced difference, with a male/female ratio of 13.69. Xanthine, a key intermediate in purine metabolism, had a male/female ratio of 7.76. The male/female ratio for L-histidinol was 5.78, and that for stearamide was 1.5 7, the latter being associated with plant allelopathy (Fig 1b 2b and 2c).

### Sex Specificity of Root Exudates in Juvenile Plants

In the juvenile group, the female/male ratio for phosphoenolpyruvate was 32.96, for pyridoxal (vitamin B₆) it was 20.78, for L-valine it was 14.18, and for ALPHA-humule ne it was 15.39. Phenylglyoxylic acid was an exudate unique to juvenile female plants (Fig 3a and 4a).

**Fig 3.**
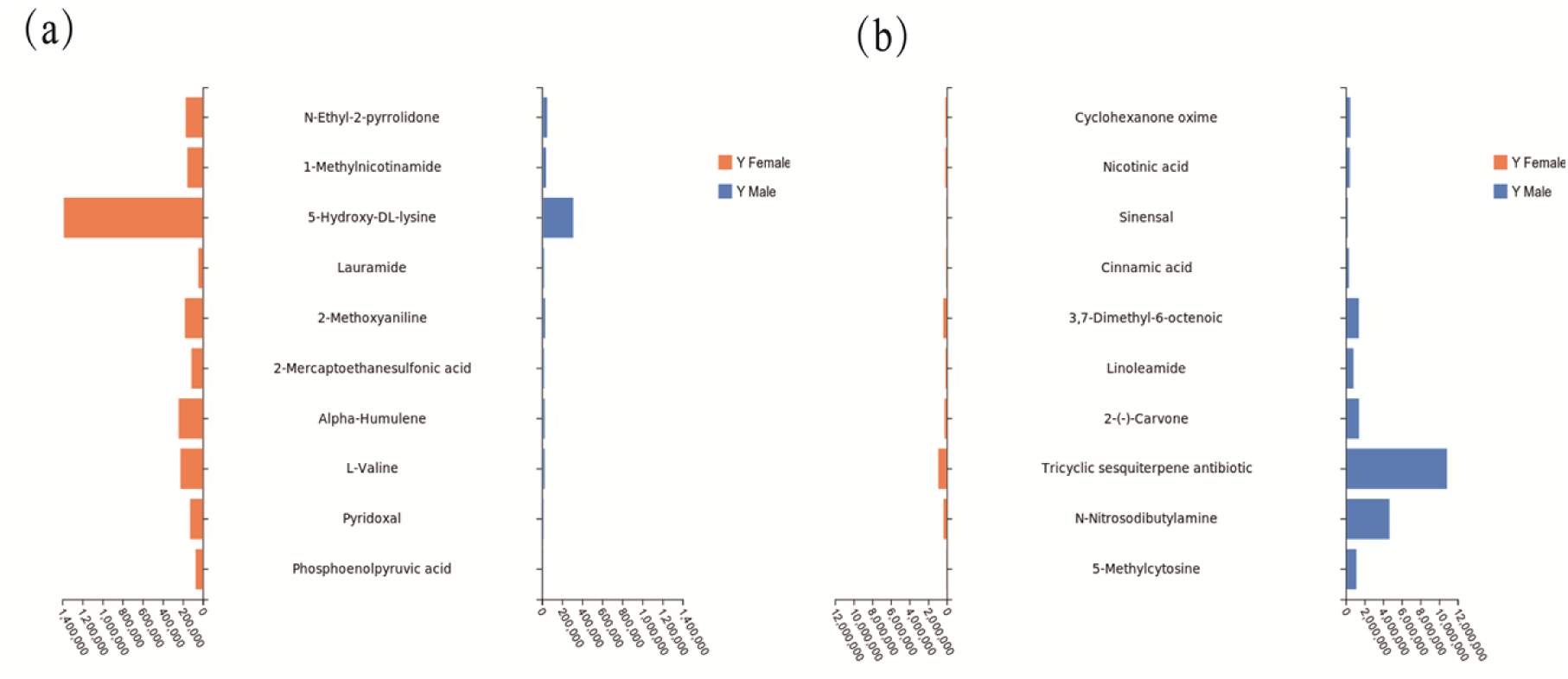
Pyramid plot of differential peak areas of root exudates in juvenile Taxus cuspidata. (a) represents the peak areas of the top 10 substances with higher abundance in juvenile female plants than in juvenile male plants. (b) represents the peak areas of the top 10 substances with higher abundance in juvenile male plants than in juvenile female plants.

**Fig 4.**
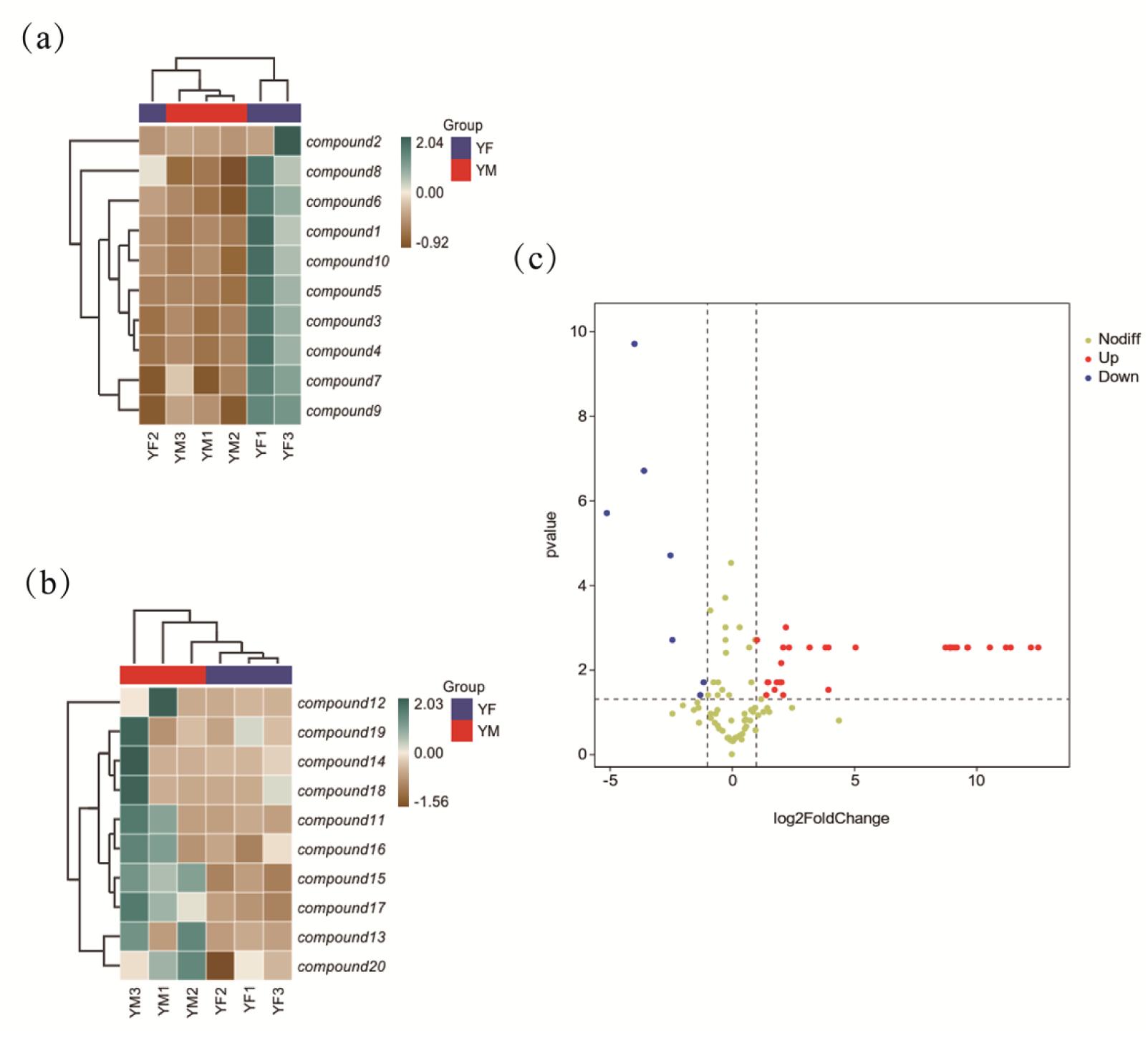
Heatmap and volcano plot of differential root exudates in juvenile Taxus cuspidata. (a) represents the heatmap of the top 10 substances with higher abundance in juvenile female plants than in juvenile male plants. (b) represents the heatmap of the top 10 substances with higher abundance in juvenile male plants than in juvenile female plants. (c) represents the volcano plot of significant differences in root exudates between juvenile male and female plants.

Among the compounds with higher peak values in juvenile male plants than in juvenile female plants, 5-methylcytosine showed a male/female ratio of 34.45, with peak values in males far exceeding those in females. N-nitrosodibutylamine had a male/female ratio of 13.16, and 2-(-)-carvone (a monoterpenoid compound) had a male/female ratio of 5.7. Tricyclic sesquiterpenoid antibiotics (male/female ratio 11.93) also exhibited significant sex differences (Fig 3b 4b and 4c).

### Differences in root exudates between adult and juvenile plants

Comparison between adult and juvenile plants revealed that the secretion intensity of adult plants was substantially higher than that of juvenile plants, often exceeding by several orders of magnitude. For example, fosfomycin exhibited the greatest difference between the two stages, with a peak value of 20.5 M in adult plants compared to only 88.74 K in juvenile plants. Cinnamic acid and hyocholic acid also showed markedly higher levels in adult plants than in juvenile plants. Furthermore, the mean peak value of phenylglyoxylic acid in adult plants reached 8.879 M, whereas it was undetectable in juvenile plants. 2-Ethylphenol was secreted at extremely high levels in adult female plants but was present at very low levels in juvenile female plants. A juvenile-specific compound, 1,3-dicyclohexylurea, was not detected in adult plants. Additional compounds with substantial differences between developmental stages are presented in Tables 2 and 3.

**Table 2.** Differences in root exudates of female *Taxus cuspidata* plants between adult and juvenile stages.

| Compound name | Adult female mean | Juvenile female mean | Fold change (Adult/Juvenile) |
| --- | --- | --- | --- |
| Fosfomycin | 20,497,129.0 | 88,741.0 | 231-fold |
| Uracil | 2,657,743.7 | 98,925.7 | 26.9-fold |
| Phosphoenolpyruvate | 1,558,416.3 | 70,318.3 | 22.2-fold |
| Bisaborusadinine | 1,234,737.3 | 77,013.7 | 16.0-fold |
| Cinnamic acid | 746,929.7 | 56,737.7 | 13.2-fold |
| Pyridoxal | 1,484,631.7 | 126,776.0 | 11.7-fold |
| Nicotinic acid (Vitamin B3) | 475,148.3 | 163,335.0 | 2.9-fold |

**Table 3.**
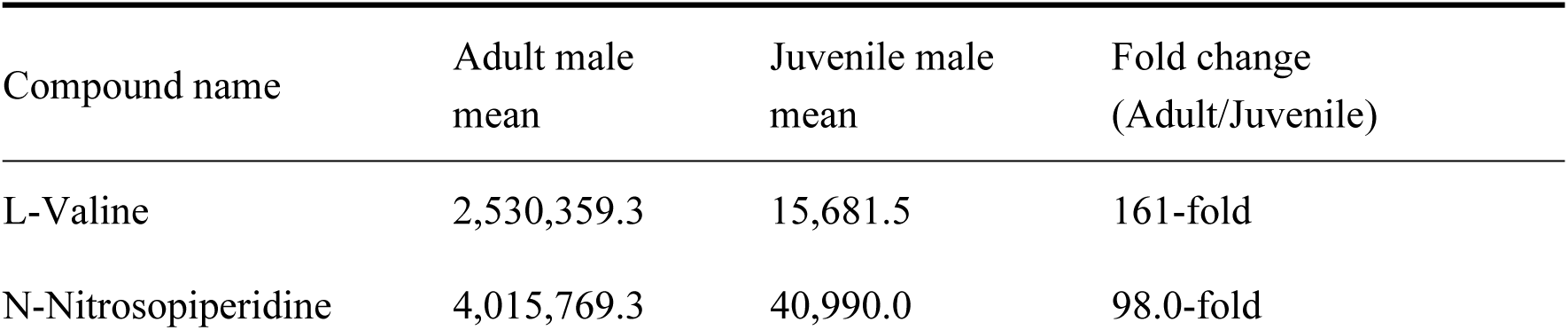
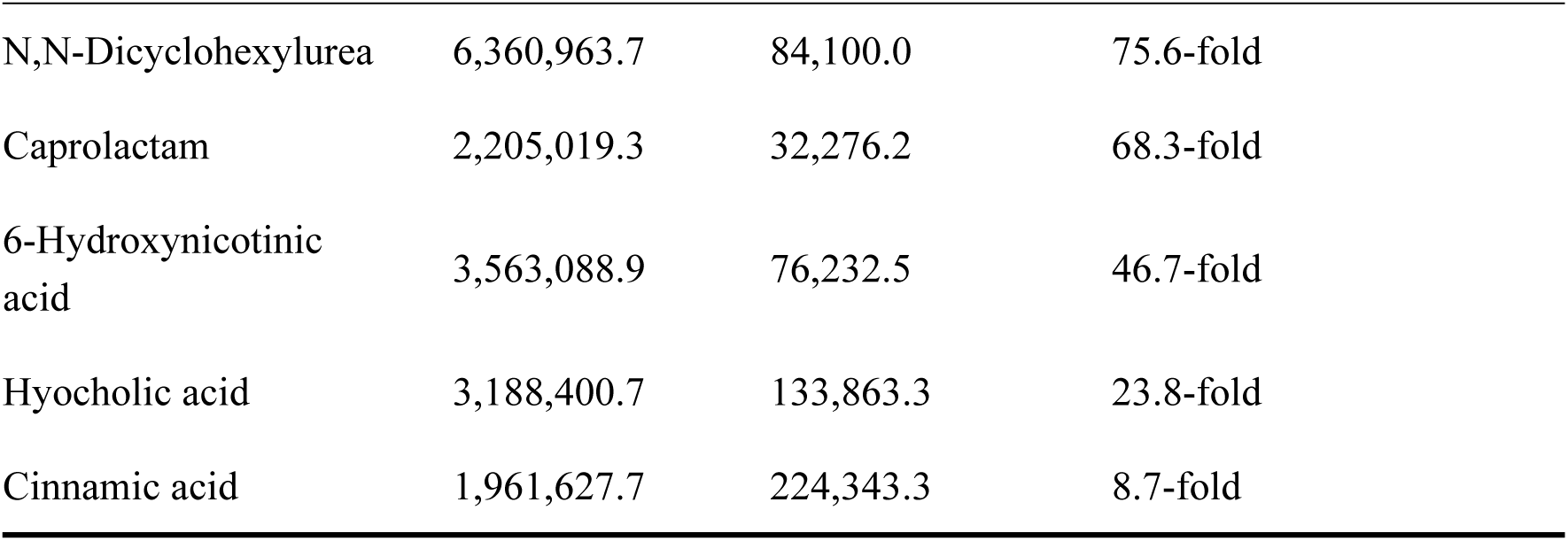
Differences in root exudates of male *Taxus cuspidata* plants between adult and juvenile stages.

### Sex-based differences in rhizosphere microbiota

#### Sex-based differences in rhizosphere microbial abundance

Resampling and normalization were performed on the rhizosphere soil microbial comm unity sequences of male and female plants. The results showed that the rarefaction abundance for bacterial samples was 25,318, with a total of 17,579 bacterial ASVs identified. For fungal samples, the rarefaction abundance was 77,095, with a total of 2,637 fungal ASVs identified.

Analysis of the rhizosphere microbial communities of male and female plants revealed significant sex specificity. Adult female rhizosphere soil contained 7,507 bacterial OTUs, while adult males had 6,199 OTUs, with 1,853 shared OTUs. Adult female rhizosphere soil contained 925 fungal OTUs, adult males had 858 OTUs, and the number of shared OTUs was 322.

The dominant microbial groups in the rhizosphere soil of juvenile plants differed from those of adult plants. Juvenile females had 5,665 bacterial OTUs, juvenile males had 6,334 OTUs, with 1,788 shared OTUs. Juvenile female rhizosphere soil contained 91 8 fungal OTUs, juvenile males had 1,046 OTUs, with 346 shared OTUs, indicating higher diversity in juvenile males (Fig 5a).

**Fig 5.**
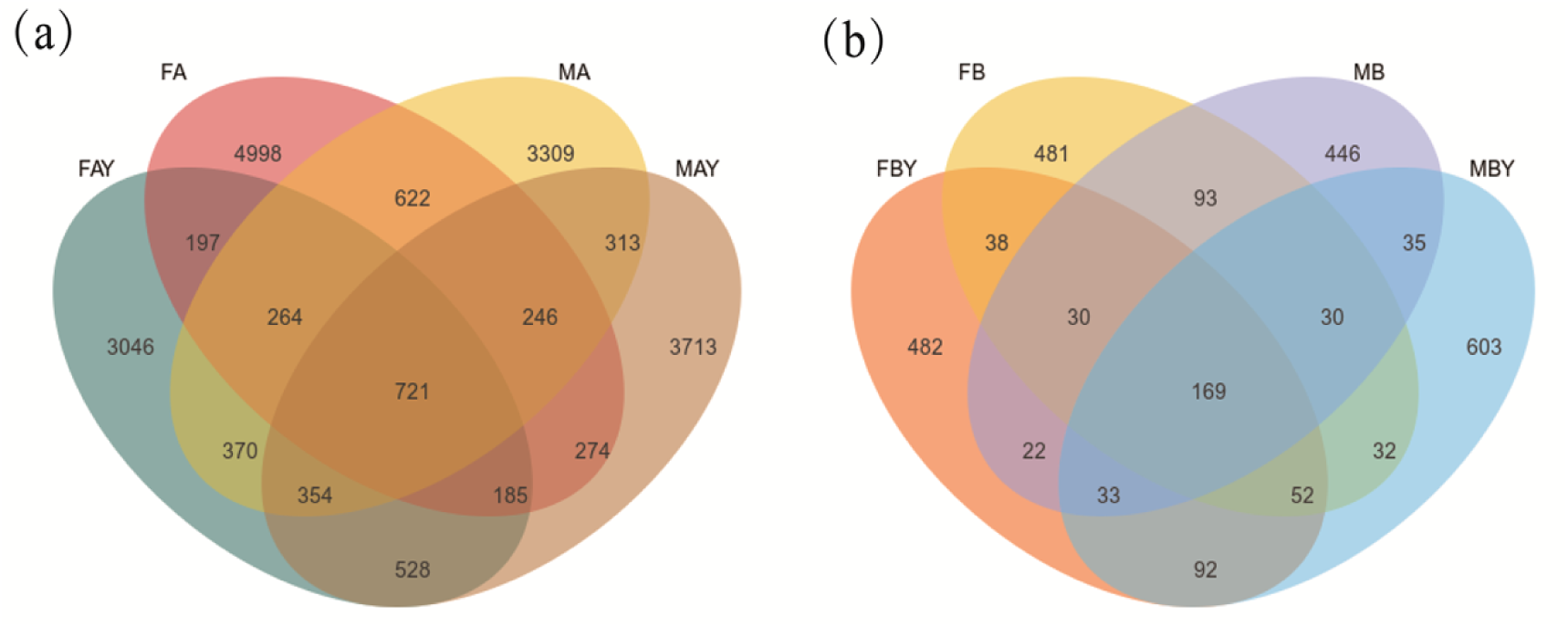
Venn diagram of differential rhizosphere microorganisms in *Taxus cuspidata.* FA represents adult female plants, FAY represents juvenile female plants. MA represents adult male plants, MAY represents juvenile male plants. (a) Venn diagram of rhizosphere soil bacteria of *Taxus cuspidata.* (b) Venn diagram of rhizosphere soil fungi of *Taxus cuspidata*.

#### Developmental stage-dependent differences in rhizosphere microbial abundance

Comparison of rhizosphere soil microorganisms between adult and juvenile plants revealed substantial differences across age stages. Adult females had 7,507 bacterial OTUs, while juvenile females had 4,298 OTUs, with 1,367 shared OTUs. Adult females had 925 fungal OTUs, juvenile females had 918 OTUs, with 289 shared OTUs, showing minimal differences in the richness of unique microbial taxa between adult and juvenile females.

Adult males had 6,199 bacterial OTUs, while juvenile males had 6,334 OTUs, with 1,634 shared OTUs. Adult males had 858 fungal OTUs, juvenile males had 1,046 OTUs, with 267 shared microbial OTUs, indicating that unique fungal taxa were more abundant in the soil of juvenile males (Fig 5b).

#### Sex specificity of bacterial species in rhizosphere soil

Sequencing identified a total of 39 bacterial phyla and 1,107 species, as well as 11 fungal phyla and 811 species, in the rhizosphere soil of *Taxus cuspidata*. Significant specificity was observed with respect to sex and developmental stage.

At the phylum level, Proteobacteria, Acidobacteriota, and Actinobacteriota were the core dominant phyla. Adult females (FA) enriched Proteobacteria (27.96%) and Gemmatimonadota, whereas adult males (MA) enriched Actinobacteriota (20.52%) and Chloroflexi. This sex difference was less pronounced at the juvenile stage; however, the abundances of Verrucomicrobiota and Myxococcota in juvenile males (MAY) were nearly twice those in juvenile females (FAY), reflecting the complex regulation of rhizosphere balance by juvenile males.

At the genus level, *RB41* (belonging to Acidobacteriota) was a signature bacterial group in adult plants, with extremely high abundance in females (2.24%), but was nearly absent in juvenile plants. *AD3* (belonging to Chloroflexi) dominated the male rhizosphere, especially in adult males (14.67%), significantly higher than in females (9.09%). In contrast, *Candidatus Solibacter* and *Acidothermus* were enriched at the juvenile stage, reflecting the adaptability of the juvenile rhizosphere environment to acidic conditions.

Across different age stages, Mortierellomycota and its genus Mortierella (8.24%) were signature differential groups in adult females, significantly higher than in juvenile females. The proportion of *Penicillium* also increased with plant growth. In juvenile females, the dominant group was Pseudopyrenula (4.81%), which was 3.84 times higher than in adult females. In adult males, *Pseudogymnoascus* (37.52%) became a core fungal group, far exceeding its proportion at the juvenile stage. Juvenile males had higher proportions of *Humicola* and *Fusarium* than adult males, with Ascomycota (90.15%) being the absolutely dominant phylum (Fig 6a and 6b).

**Fig 6.**
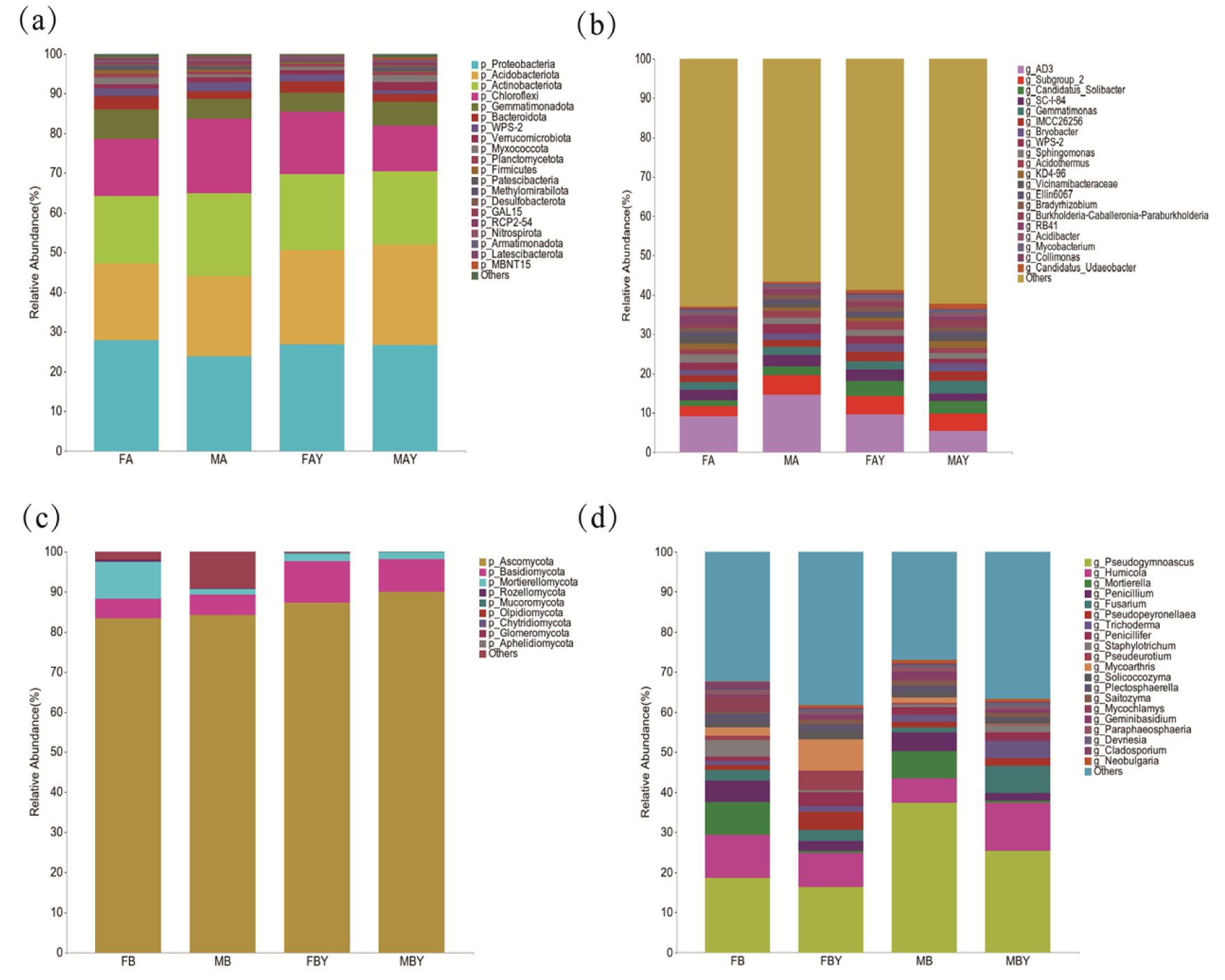
Bar plot of differential rhizosphere microorganisms in Taxus cuspidata. (a) Specificity of rhizosphere soil bacteria at the phylum level. (b) Specificity of rhizosphere soil bacteria at the genus level. (c) Sex specificity of rhizosphere soil fungi at the phylum level. (d) Specificity of rhizosphere soil fungi at the genus level.

#### Specificity of fungal species in rhizosphere soil

At the fungal phylum level, Ascomycota (over 80%) was absolutely dominant across all rhizosphere soil samples. The most significantly differential fungal group was Mortierellomycota. The abundance in adult females (FB) (9.13%) was 6.4 times that in adult males (MB) and much higher than that at the juvenile stage, making it a characteristic phylum of adult females. Additionally, the abundance of Basidiomycota was generally higher at the juvenile stage (FBY/MBY) than at the adult stage.

At the genus level, *Pseudogymnoascus* was the dominant group in the male rhizosphere, accounting for as much as 37.52% in adult males (MB), significantly higher than in females. The female rhizosphere recruited a greater diversity of functional fungal genera, such as *Humicola*, *Mortierella*, and *Mycochlamys*.

Across different growth and developmental stages, juvenile females (FBY) enriched *Pseudeurotium* at a level 43.6 times that of males. As plants developed, the proportion of unclassified fungi increased significantly in adult males (9.11%), indicating the presence of more unknown differences in the rhizosphere environment of *Taxus cuspidata* after reaching adulthood (Fig 6c and 6d).

#### Rhizosphere Soil Microbial α-Diversity Analysis

In the adult stage, female plants exhibited significantly higher Chao1 index (2943.22) and Shannon index (11.35) compared to male plants (Chao1: 2390, Shannon: 9.87), indicating that adult females possess greater bacterial richness and diversity.

In the juvenile stage, male plants showed significantly higher Chao1 index (2670) and Shannon index (10.33) than female plants, suggesting that juvenile male rhizospheres recruit more abundant bacterial species. Differences in the Simpson index among groups were small, with no absolutely dominant bacterial species (Fig 7a and 7c).

**Fig 7.**
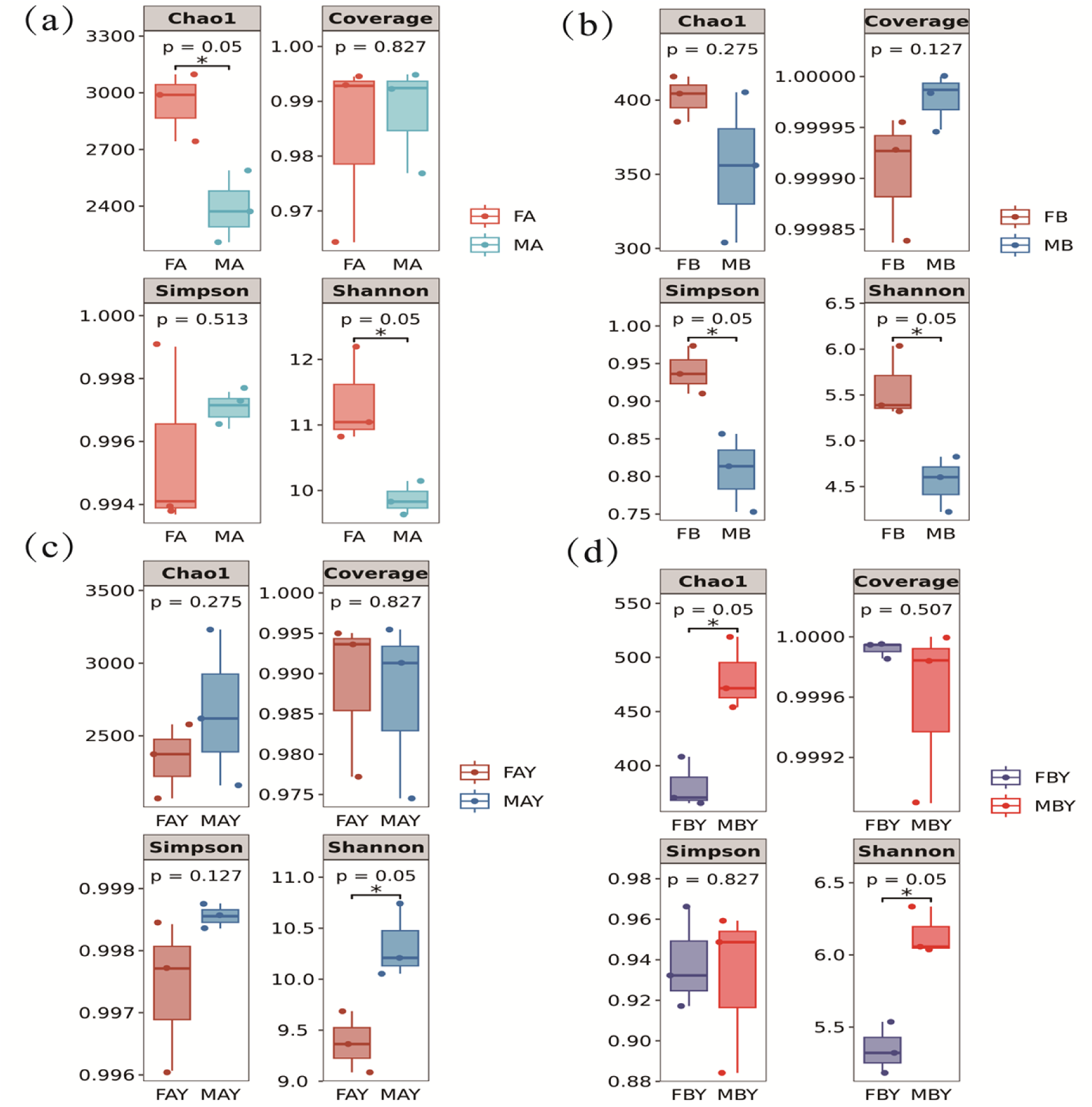
Differential α-diversity of rhizosphere microorganisms in Taxus cuspidata. (a) represents bacterial α-diversity in the rhizosphere soil of adult plants. (b) represents fungal α-diversity in the rhizosphere soil of juvenile plants. (c) represents fungal α-diversity in the rhizosphere soil of adult plants. (d) represents fungal α-diversity in the rhizosphere soil of juvenile plants.

The α-diversity trend of fungal communities was generally consistent with that of bacteria, but the differences between sexes were more pronounced. In the adult stage, female plants had significantly higher Chao1 index and Shannon index (5.58) than male plants (4.55). The Simpson index of the MB group (0.81) was significantly lower than that of the FB group (0.96), indicating that the dominant fungal groups in the adult male rhizosphere were highly concentrated, with lower community evenness.

In the juvenile stage, juvenile males exhibited stronger community dominance, with the Chao1 index (481.54) being 26.3% higher than that of juvenile females (381.38), and a higher Shannon index (males 6.14 vs females 5.34), indicating that the rhizosphere fungal community structure of juvenile males is more complex and stable.

Comparison of data across different developmental stages revealed that from juvenile to adult stages, fungal richness in females increased steadily (FB > FBY), whereas fungal diversity in males fluctuated dramatically. The Chao1 index of adult males (MB) (355) significantly decreased compared to their juvenile stage (481.54), and the community distribution became highly uneven.

Adult females possess higher and more stable microbial community diversity, whereas males exhibit stronger microbial recruitment ability during the juvenile stage. This shift in diversity patterns is closely related to the resource allocation strategies and metabolite changes of different sexes during growth and development (Figs 7b and 7d).

#### Rhizosphere Soil Microbial β-Diversity Analysis

In the NMDS (non-metric multidimensional scaling) plots, each point represents a biological sample, with different colors used to distinguish different sample groups, such as adult male/female plants and juvenile male/female plants. The distance between samp les approximately reflects the similarity of community composition: the closer the points, the more similar the communities, and the greater the distance, the larger the difference (Fig 8a and 8b).

**Fig 8.**
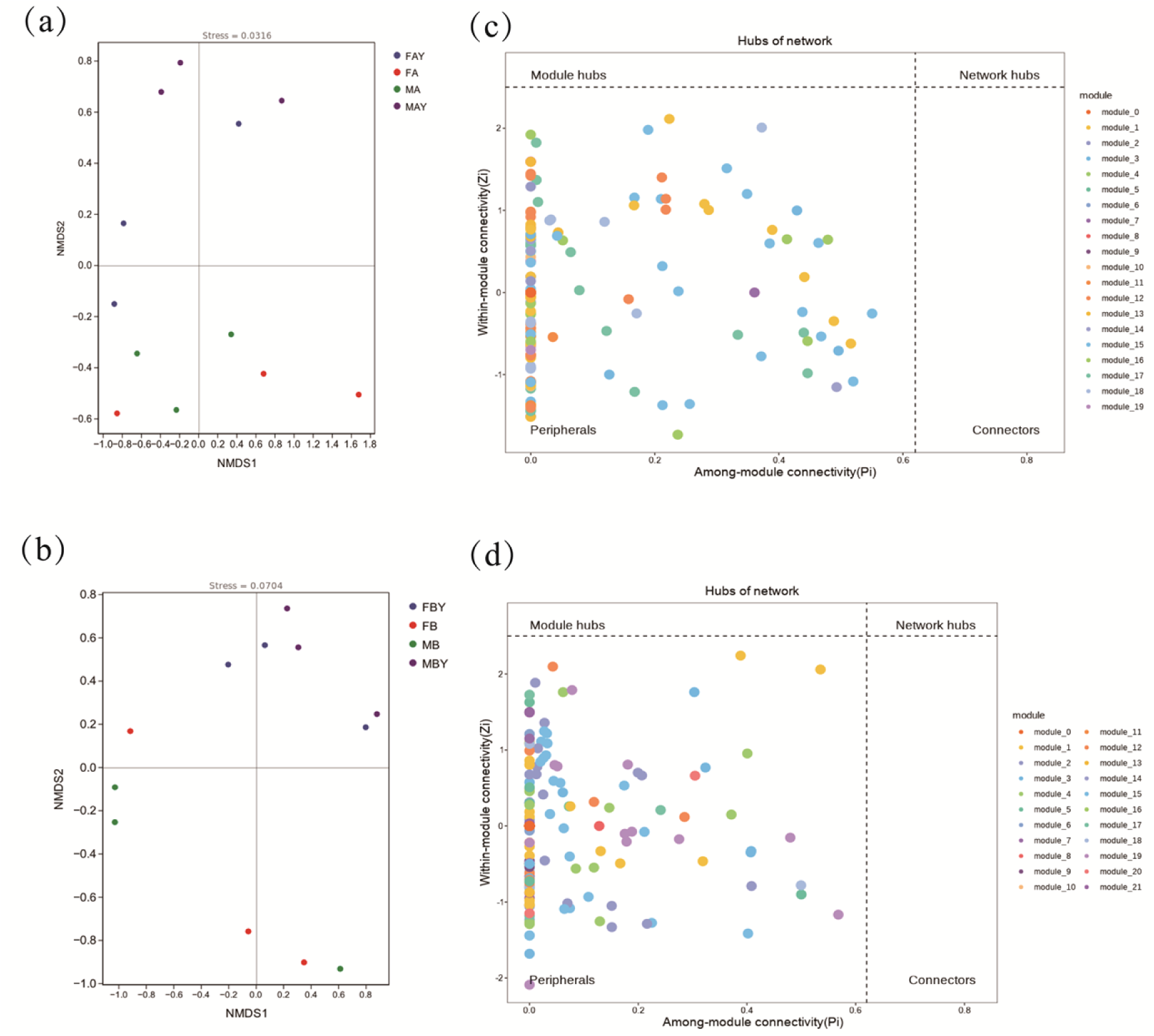
Differential β-Diversity and network diagram of rhizosphere microorganisms in Taxus cuspidata. (a) represents bacterial β-diversity in the rhizosphere soil. (b) represents fungal β-diversity in the rhizosphere soil. (c) represents the hub-node diagram of rhizosphere soil bacteria. (d) represents the hub-node diagram of rhizosphere soil fungi.

#### Analysis of Rhizosphere Microbial Network

Analysis of 200 ASVs in the bacterial community using a hub-node diagram revealed that all 200 ASV nodes were classified as “Peripherals” (peripheral nodes), with no hub nodes or connector nodes. Only a few nodes served as inter-module connectors, while the majority of nodes participated in local intra-module interactions. The loss of a single node has a minor impact on the overall structure, and the loose connections result in low functional synergy efficiency within the community. Both positive and negative association strengths were weak, with no strongly antagonistic or strongly synergistic interactions, indicating that the community is in a relatively stable weak equilibrium state (Fig 8c).

Network analysis of 158 fungal ASV nodes showed that all nodes were “Peripherals” (peripheral nodes), with no hub nodes or connector nodes, consistent with the characteristics of the soil bacterial network but with weaker interaction strengths. The network exhibited strong resistance to disturbance, yet also extremely low synergy efficiency, relying on weak interactions to maintain community stability. Only 39 nodes served as bridges between modules, indicating thorough functional differentiation and adaptation to the heterogeneity of the belowground soil environment. The efficiency of material and signal transfer in the fungal community is low, reflecting a state of weak synergy and weak competition equilibrium (Fig 8d).

#### Correlation analysis between root exudates and rhizosphere microorganisms

Correlation analysis revealed that in female samples, the abundances of uracil and pyridoxal-5-phosphate were significantly higher, simultaneously attracting *Penicillium* and *Mortierella*. 6-Hydroxynicotinic acid was extremely abundant in males and showed a strong positive correlation with *Pseudogymnoascus*. Oleamide was enriched in female plants, whereas stearamide was enriched in male plants.

In juvenile male plants, the peak areas of 5-methylcytosine and tricyclic sesquiterpenoid antibiotics were far higher than those in juvenile females, and they were strongly correlated with *Fusarium*. L-Valine and pyridoxal were highly enriched in females, and *Pseudogymnoascus* exhibited a positive correlation trend with nicotinic acid (vitamin B_3_)(Table 4,Table 5).

**Table 4.** Predictive correlation analysis between root exudates and rhizosphere microorganisms of adult Taxus chinensis.

| Metabolite | Microbial Genus | Correlation Coefficient (ρ) | P-value |
| --- | --- | --- | --- |
| 6-Hydroxynicotinic acid | *Pseudogymnoascus* | 0.943 | 0.005** |
| Uracil | *Staphylotrichum* | 0.886 | 0.019* |
| Xanthine | *AD3* | 0.829 | 0.042* |
| Nicotinic acid | *Mortierella* | 0.771 | 0.072 |
| Stearamide | *Subgroup_2* | 0.841 | 0.036* |

| Metabolite | Microbial Genus | Correlation Coefficient<br>($\rho$) | *P*-value |
| --- | --- | --- | --- |
| 9-Methyl-3-tropinone | *Mycochlamys* | 0.928 | 0.008** |

**Table 5.** Predictive correlation analysis between root exudates and rhizosphere microorganisms of juvenile Taxus chinensis.

| Metabolite | Microbial Genus | Correlation Coefficient<br>($\rho$) | *P*-value |
| --- | --- | --- | --- |
| 5-Methylcytosine | *Fusarium* | 0.943 | 0.005** |
| Tricyclic sesquiterpene antibiotic | *Trichoderma* | 0.886 | 0.019* |
| L-Valine | *Candidatus_Solibacter* | 0.829 | 0.042* |
| Nicotinic acid | *Pseudogymnoascus* | 0.771 | 0.072 |
| 2-(-)-Carvone | *Penicillifer* | 0.812 | 0.049* |

## Discussion

According to the parental investment theory proposed by Lloyd and colleagues, female plants invest more energy and mineral nutrients than male plants during reproduction (flowering and fruiting) for ovule development and seed production[32–33]. In contrast, male plants allocate most of their resources to the vegetative growth of aboveground leaves and branches to enhance photosynthetic nutrient accumulation, future reproductive competition, and self-defense[34–35]. This resource allocation strategy leads to significant differences in the types, age-related patterns, and abundances of root exudates between sexes, thereby influencing belowground chemical signaling and biological interaction patterns.

### Sexual and Developmental Stage-Specific Differences in Root Exudates

Dominant role of developmental stage: For example, root exudates of Arabidopsis thaliana are predominantly composed of sugars at early stages, whereas amino acids and phenolic compounds increase in exudates from 30-day-old senescing plants[36]. Studies on maize have demonstrated that plant developmental stage is a major determinant of root exudate composition and quantity; carbon exudation per plant increases over time, while the carbon exudation rate per unit surface area decreases as plants mature[37]. The metabolite profiles of oat (*Avena sativa*) at early growth stages differ considerably from those at mid- and late developmental stages[8]. In the present study, root exudates of *Taxus cuspidata* exhibited distinct characteristics at different developmental stages. Young female plants secreted high concentrations of phosphoenolpyruvate (PEP) and L-valine, confirming that they are in a critical period of energy metabolism and protein synthesis. Additionally, they constructed a multi-component defense system comprising tropinone, β-caryophyllene, and other compounds. Young male plants secreted 5-methylcytosine and tricyclic sesquiterpenoid antibiotics to enhance their competitive ability and defense capacity.

Dominant role of sexual specificity: Previous studies have shown that female *Populus cathayana* releases higher concentrations and more diverse phenolic allelochemicals into the soil compared to males[20]. At the adult stage, female exudates are enriched with antioxidants (e.g., 2-mercaptoethanesulfonic acid) and methionine metabolism-related substances, thereby enhancing the antioxidant defense system to cope with the stress induced by reproductive investment[38]. In contrast, adult male plants secrete higher amounts of alpha-humulene (a terpenoid allelochemical) and xanthine, favoring niche competition and defense against adverse environments through chemical means[39]. By utilizing volatile terpenoids to inhibit other plants, male plants secure their own resource acquisition.

### Sexual and Developmental Stage-Specific Differences in Rhizosphere Microorganisms

Root exudates are the primary factors shaping the rhizosphere microbial community, providing essential carbon sources and energy for rhizosphere microorganisms and driving their metabolic activities[8]. The establishment of the rhizosphere microbial community is a dynamic process, and its composition changes with plant growth. A study on Arabidopsis alpina revealed three distinct rhizosphere microbial communities over a 28 -week growth period[40]. Similar findings have been reported in potato, where β-proteobacteria involved in nutrient cycling significantly increase during the late growth stage[41]. This study supports the “host-mediated selection” theory[42], revealing how *Tax us cuspidata* recruits’ sex-specific microorganisms through differential root exudation. According to the “cry for help” hypothesis proposed by Rudrappa et al., plants alter their root exudates under stress to recruit beneficial microorganisms[43]. Young male plants recruit a large number of stress-tolerance-associated microbiota, including Actinobacteriota and *Flectobacillus*, by secreting tricyclic sesquiterpenoid antibiotics, thereby establishing a growth advantage during early population competition. Young female plants, in contrast, prioritize microbial screening and reservation by recruiting genera such as *Pseudogymnoascus* and *Pseudopeyronellaea*, allocating more resources to their own growth and energy accumulation in preparation for subsequent reproductive processes such as flowering and fruiting.

Adult female plants tend to recruit microorganisms with nutrient-conversion capabilities, compensating for the nutrients consumed by reproduction through improved soil nutrient utilization efficiency[44]. Specifically, Proteobacteria and *Mortierella* are the dominant taxa in female plants, whereas Actinobacteriota and *Flectobacillus* account for a higher proportion in male plants. Similar phenomena have been observed in studies on Hippophae tibetana and Ginkgo biloba[45–46]. These findings indicate that female plants recruit microorganisms more for nutrient acquisition, whereas male plants recruit microorganisms more for environmental defense.

### Recruitment of Rhizosphere Microorganisms by Sex-Specific Root Exudates

Plants and microorganisms do not exist as loosely assembled entities but rather as co-evolved units, in which root exudates play an important role[47]. Root exudates can increase carbon composition and provide energy for microorganisms, thereby significantly enhancing microbial biomass[1]. By secreting specific root exudates (e.g., phenolic acids, sugars, flavonoids, and organic acids), plants can selectively recruit microorganisms that are beneficial to their growth[48–51].

Correlation analysis (Tables 4 and 5) and functional association predictions indicate that phenolic compounds secreted by female *Taxus cuspidata* plants (e.g., 5-methylcytosine and 2-ethylphenol) are significantly positively correlated with Cystofilobasidium and Fusarium. These microorganisms participate in the synthesis of secondary metabolites an d nutrient transformation in female plants, synergistically supporting their reproductive cycle.

Male defense recruitment module: The tricyclic sesquiterpenoid antibiotics secreted by male plants are highly correlated with pathogen-inhibiting microorganisms and Pseudogymnoascus. This mechanism not only defends against pathogens but also enhances soil fertility through Pseudogymnoascus (which decomposes complex compounds) by degrading complex soil organic matter, thereby meeting the pollen production demands of m ale plants.

Just as Arabidopsis thaliana releases malic acid to attract *Bacillus subtilis*[43,52], and the diverse substances in root exudates of Tangut grape (*Vitis tangutica*) are closely related to the richness of soil microbial species[29], root exudates ensure that plants of different sexes can recruit specific functional microbial groups at key developmental stages (e.g., male flowering and female fruiting).

This study provides a new perspective for belowground ecology research on dioecious plants and demonstrates that understanding the composition and function of rhizosphere microbial communities associated with dioecious plants is of great significance for revealing evolutionary differences between female and male plants. In practical applications, sexual-specific probiotics or functional cultivation substrates can be developed based on the preference of female and male seedlings for root exudates, thereby significantly enhancing plant stress resistance and survival rates. Furthermore, the study reveals that male plants have a stronger ability to recruit microbial communities under stressful conditions. Therefore, it is recommended to prioritize the use of male plants in stressful environments to improve the stability of artificial forests.

## Conclusion

The root exudates of *Taxus cuspidata* exhibit significant sex specificity and developmental stage differences. Adult female plants tend to enrich antioxidant and nutrient metabolism substances to support reproductive investment, whereas male plants secrete mor e terpenoid allelochemicals to enhance survival competition. At the juvenile stage, fem ale exudates contain higher concentrations of phosphoenolpyruvate (PEP) and L-valine, storing nutrients for future reproduction. In contrast, juvenile male plants secrete 5-m ethylcytosine and tricyclic sesquiterpenoid antibiotics, enhancing their own competitive and defensive capabilities.

The composition of rhizosphere microorganisms in *Taxus cuspidata* shows clear sex-related differences. Female plants mainly enrich growth-promoting bacteria, while male plants recruit more decomposer fungi, both of which are significantly regulated by root exudates.

Rhizosphere microbial diversity exhibits a “sex reversal” phenomenon across developmental stages: microbial richness is higher in female plants during the adult stage, whereas juvenile males recruit a more diverse microbial community. These results indicate that dioecious plants regulate rhizosphere microbial communities through differential root exudates, thereby creating a rhizosphere environment co-shaped by sex and developmental stage.

## Notes

**Competing interest statement:** The authors declare that they have no known competing financial interests or personal relationships that could have appeared to influence the work reported in this paper.

### Competing Interest Statement

The authors have declared no competing interest.

